# The small noncoding RNA sr8384 determines solvent synthesis and cell growth in industrial solventogenic clostridia

**DOI:** 10.1101/663880

**Authors:** Yunpeng Yang, Nannan Lang, Huan Zhang, Lu Zhang, Changsheng Chai, Weihong Jiang, Yang Gu

## Abstract

Small noncoding RNAs (sncRNAs) are crucial regulatory molecules in organisms and are well known not only for their roles in the control of diverse essential biological processes but also for their value in genetic modification. However, to date, in gram-positive anaerobic solventogenic clostridia (which are a group of important industrial bacteria with exceptional substrate and product diversity), sncRNAs remain minimally explored, leading to a lack of detailed understanding regarding these important molecules and their use as targets for genetic improvement. Here, we performed large-scale phenotypic screens of a transposon-mediated mutant library of *Clostridium acetobutylicum*, a typical solventogenic clostridial species, and discovered a novel sncRNA (sr8384) that functions as a determinant positive regulator of growth and solvent synthesis. Comparative transcriptomic data combined with genetic and biochemical analyses revealed that sr8384 acts as a pleiotropic regulator and controls multiple targets that are associated with crucial biological processes, through direct or indirect interactions. Notably, modulation of the expression level of either sr8384 or its core target genes significantly increased the growth rate, solvent titer and productivity of the cells, indicating the importance of sr8384-mediated regulatory network in *C. acetobutylicum*. Furthermore, a homolog of sr8384 was discovered and proven to be functional in another important *Clostridium* species, *C. beijerinckii*, suggesting the potential broad role of this sncRNA in clostridia. Our work showcases a previously unknown potent and complex role of sncRNAs in clostridia, providing new opportunities for understanding and engineering these anaerobes, including pathogenic *Clostridium* species.

**IMPORTANCE:** The discovery of sncRNAs as new resources for functional studies and strain modifications are promising strategies in microorganisms. However, these crucial regulatory molecules have hardly been explored in industrially important solventogenic clostridia. Here, we identified sr8384 as a novel determinant sncRNA controlling cellular performance of solventogenic *Clostridium acetobutylicum* and performed detailed functional analysis, which is the most in-depth study of sncRNAs in clostridia to date. We reveal the pleiotropic function of sr8384 and its multiple direct and indirect crucial targets, which represents a valuable source for understanding and optimizing this anaerobe. Of note, manipulation of these targets leads to improved cell growth and solvent synthesis. Our findings provide a new perspective for future studies on regulatory sncRNAs in clostridia.

## INTRODUCTION

Historically, the application of solventogenic clostridia in the large-scale production of the bulk chemicals acetone, n-butanol and ethanol, a process called ABE fermentation, has demonstrated the value of these anaerobic microorganisms (1, 2). In recent years, in view of the exceptional substrate and product diversity of solventogenic clostridia, the biological production of cost-effective bulk chemicals and biofuels using *Clostridium* species as chassis has attracted renewed attention (3). To unlock the full potential of solventogenic clostridia in industrial applications, a detailed understanding of metabolic regulation and discovery of more crucial regulatory elements in these anaerobes are necessary. However, to date, this aspect remains minimally explored, and only a limited number of transcription factors from solventogenic clostridia have been identified and subjected to functional analysis (4); in addition, other types of regulatory molecules and modes (e.g., post-transcriptional and post-translational modes) remain largely unexplored. The lack of knowledge regarding these aspects will inevitably increase the difficulty in identifying new targets for strain improvement.

Small noncoding RNAs (sncRNAs) are crucial regulatory molecules in organisms (5, 6). In addition, sncRNAs have been increasingly regarded as promising targets for genetic improvement (7–9). Despite the increasing interest in the function of small RNAs in solventogenic clostridia, they remain largely unexplored in these anaerobes. To date, only a few small RNAs have been identified in solventogenic clostridia (10, 11). A newly reported regulator of SolB in *Clostridium acetobutylicum* was found to specifically regulate the expression of the genes in the *sol* locus, leading to a solvent-deficient phenotype after overexpression (12). Notably, a comprehensive list of sRNAs in 21 clostridial genomes (including two industrial *Clostridium* strains: *C. acetobutylicum* and *Clostridium beijerinckii*) has been computationally predicted, revealing a large number of sncRNAs in the genus *Clostridium* (13). This work, despite not focusing on functional analysis, strongly supports a continued investigation of the important roles of sncRNAs in industrial clostridia.

Here, we report the discovery of a novel sncRNA (sr8384) in *C. acetobutylicum*, a representative species of industrial solventogenic clostridia, based on phenotypic screening of a previously established transposon-based random mutant library (14). The sncRNA sr8384 was not identified in the previous systematic screening of the intergenic regions of *C. acetobutylicum* via computational analysis (13), indicating that this sncRNA has unique features that are distinct from those of the reported bacterial small RNAs. A series of genetic and biochemical analyses were carried out for a detailed functional analysis of sr8384, revealing a regulatory network that controls crucial phenotypes of *C. acetobutylicum*. Manipulation of sr8384 or its gene targets could effectively promote growth and solvent production, demonstrating the importance of this sncRNA as well as the related gene network in genetic improvement. Furthermore, we also identified a functional sr8384 homolog in *C. beijerinckii*, another important *Clostridium* species that is widely used in the fermentation of lignocellulose hydrolysates, indicating the important functions and broad role of sr8384-like sncRNAs in solventogenic clostridia

## RESULTS

### Phenotypic screens reveal a transposon mutant with greatly changed solvents production

In a previous study, we established a *mariner*-based transposon system in *C. acetobutylicum* ATCC 824, which generated a mutant library (more than 30,000 mutants) with high randomness (14). As a continuation of this work, we recently used this library to screen for mutants with phenotypic changes in essential traits, such as growth and solvent synthesis. According to the process shown in Figure 1A, more than 600 mutants were tested, and we obtained a transposon mutant (Tn mutant) that exhibited greatly impaired solvent formation during fermentation using glucose as the carbon source. This mutant could produce only 6.9 g/L of total solvents (acetone, butanol and ethanol) after 96 h of fermentation (Figure 1B), which is far less than the level produced by the wild-type strain, indicating the presence of a transposon insertion at an essential chromosomal position in the Tn mutant. By sequencing the reverse PCR product of the Tn mutant, we found that this mutant contained a transposon insertion in a 198-nt gene of unknown function (CAC2384) (Figure 1C).

**FIG 1.**
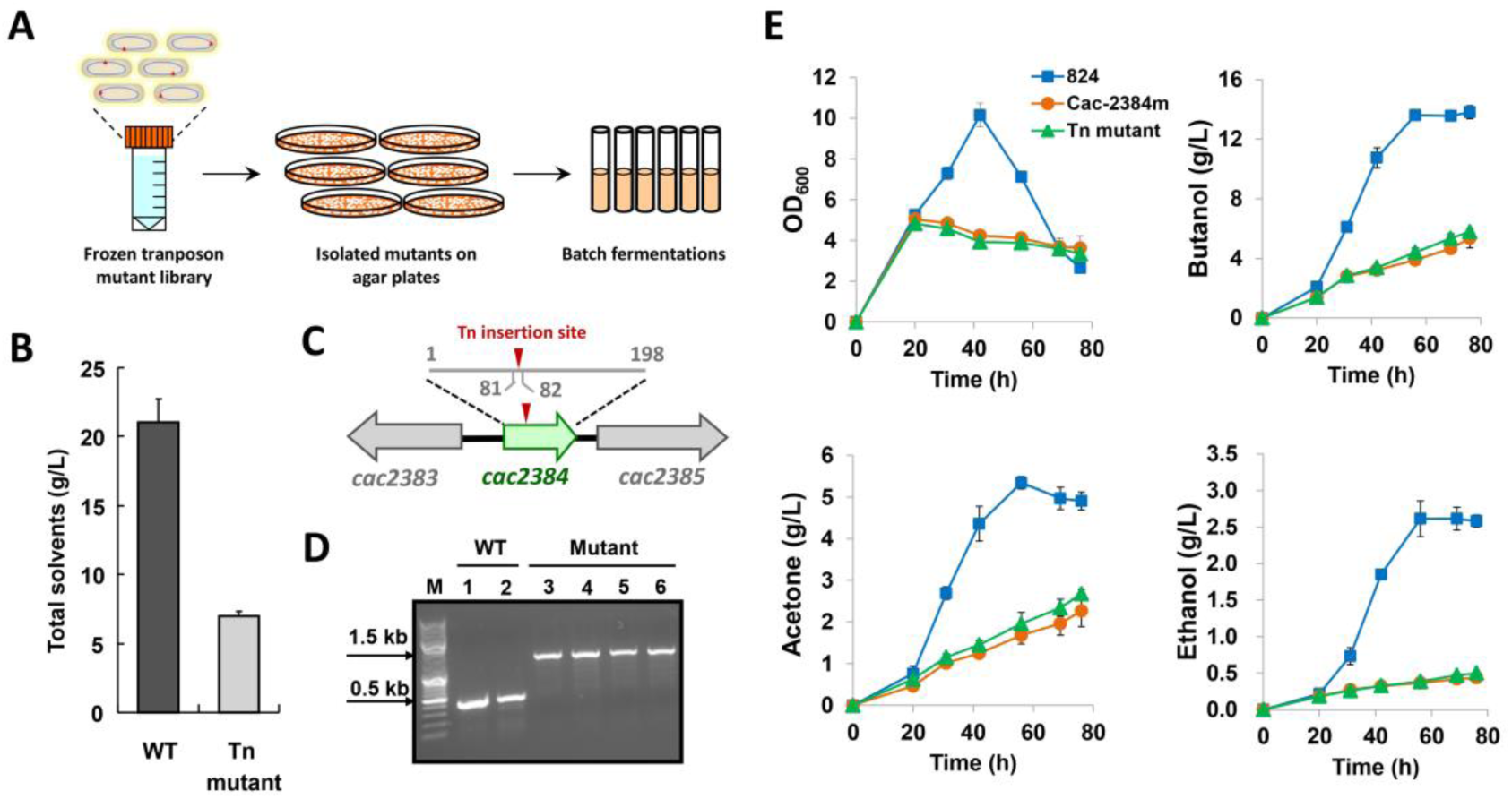
Identification and characterization of a *C. acetobutylicum* mutant with significant changes in growth and solvents production. (A) Isolation of transposon mutant with obviously altered ability in forming ABE solvents. (B) Comparison of solvents production of Tn mutant and wild-type strain. (C) Transposon insertion site (the inverted red triangle) on the chromosome of Tn mutant. It is between the +81 and +82 site of open reading frame. (D) Verification of the intron insertion in the CAC2384 gene by PCR analysis. The 1.5- and 0.5-kb band represents the PCR-amplified fragment containing the intron-inserted and original CAC2384 gene, respectively. (E) Growth and solvents formation of the Cac-2384m and wild-type strain. Data are means±standard deviations calculated from triplicate independent experiments.

### Characterization of the Tn mutant reveals a novel small noncoding RNA: sr8384

To verify whether the abovementioned phenotypic changes of the Tn mutant were due to CAC2384 inactivation, we used the group II intron-based gene inactivation method (15) to disrupt CAC2384 in wild-type *C. acetobutylicum* (Figure 1D), and the mutant obtained (named Cac-2384m) was used for phenotypic investigation. Additionally, southern blot analysis was performed to verify that the intron was incorporated only once into the genome of Cac-2384m with no other non-specific insertions (Figure S1). As expected, the Cac-2384m strain exhibited very similar profile of growth and solvent production (acetone, butanol and ethanol) to the Tn mutant strain (Figure 1E).

However, when genetic complementation was performed by separately introducing four plasmids (the plasmid pP*_2384_*-2384, which expressed CAC2384 under the control of the native promoter P*_2384_* of CAC2384; the plasmid pP*_2384_*, which harbored only the promoter P*_2384_*; the plasmid pP*_thl_*-2384, which expressed CAC2384 under the control of the constitutive promoter P*_thl_*; and the plasmid pP*_thl_*, which harbored only the promoter P*_thl_*) back into the Cac-2384m strain (Figure 2A), a surprising but interesting result was obtained: both the pP*_2384_*-2384 and pP*_2384_* plasmids could complement the deficiency of the Cac-2384m mutant in solvent formation, whereas both the pP*_thl_*-2384 and pP*_thl_* plasmids failed to do so (Figure 2B). To further convincing this finding, we performed genetic complementation experiment again through chromosomal insertion of the target DNA sequence by using the “Clostron” technology (16, 17). In brief, the abovementioned four DNA fragments were separately integrated into an intron sequence, and then inserted into the chromosome to see if the impaired phenotypes of the Cac-2384m mutant could be restored (Figure S2A, B and C). As expected, the results (Figure S2D) were similar with the abovementioned genetic complementation experiment using multicopy plasmids (Figure 2). Obviously, all these data strongly suggest that the phenotypic changes in the Cac-2384m mutant can be recovered by the independent expression of the upstream noncoding sequence (P*_2384_*) of CAC2384. In other words, there may be some crucial DNA elements in the upstream region of CAC2384, although it is unclear why the insertional disruption of CAC2384 influenced this non-coding region.

**FIG 2.**
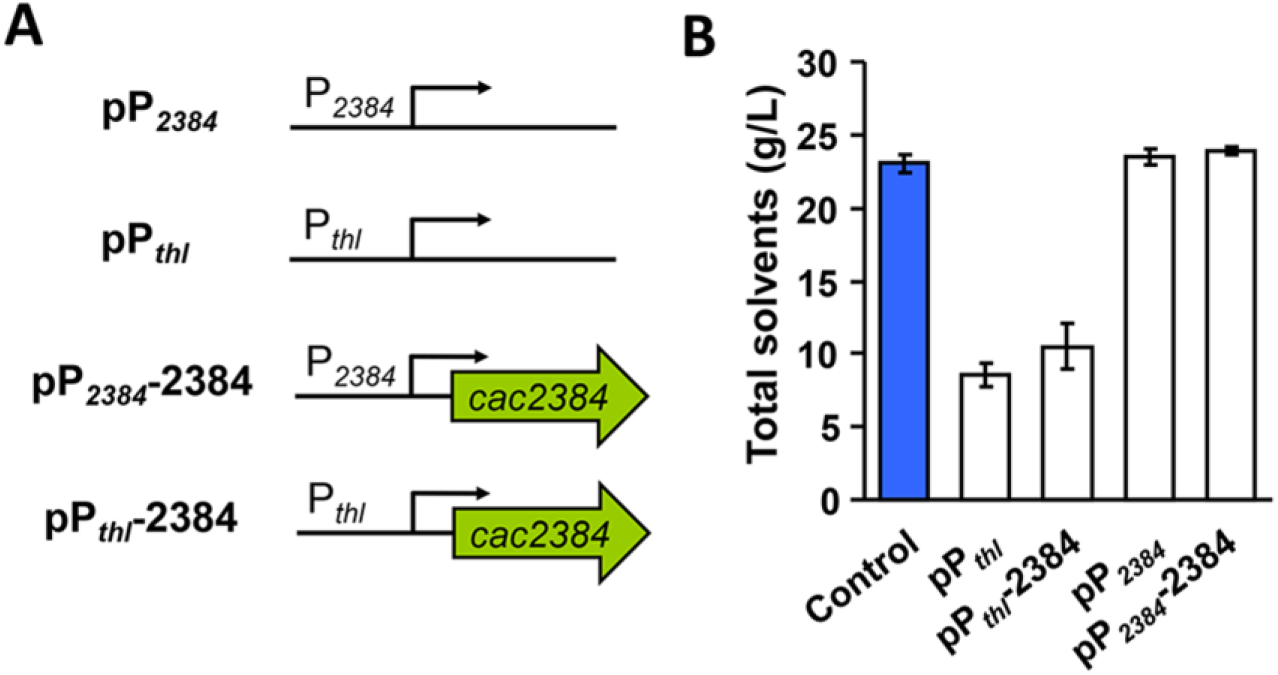
Genetic complementation of the Cac-2384m mutant indicates an unknown crucial molecule related to the phenotypic changes. (A) The four plasmids (pP*_thl_*, pP*_thl_*-2384, pP*_2384_*, pP*_2384_*-2384) constructed for genetic complementation of Cac-2384m. (B) The solvents formation of Cac-2384m mutants with the four complementary plasmids and the wild-type strain carrying an empty plasmid (Control). Data are means±standard deviations calculated from triplicate independent experiments

Next, a detailed functional analysis of the P*_2384_* sequence was performed to explore the above hypothesis. The whole sequence (202 nt) of P*_2384_* was gradually truncated, yielding 10 truncated fragments, i.e., P*_2384_* minus 10, 20, 30, 40, 50, 60, 100, 120, 140 or 150 nt. These DNA fragments were integrated into the expression plasmid and then introduced into the Cac-2384m mutant for genetic complementation analysis (Figure S3A). The results showed that all the truncated fragments retained complementation functions except the shortest fragment (with a 150-bp deletion) (Figure S3B), thus suggesting that the potential DNA element suggested above is located within the 62-nt P*_2384_*_-140_ sequence. Given the very low chance that this short 62-nt sequence encodes a functional protein, we reasoned that it may encode a small RNA (sRNA).

To explore this possibility, the following experiments were performed in sequence: (i) a two-step RT-PCR analysis for determining the transcriptional direction of the P*_2384_*_-140_ sequence; RACE (5’ and 3’ rapid amplification of cDNA ends) experiment aiming to determine the actual transcript length of P*_2384_*_-140_; (ii) Northern blotting to verify the role of this transcript (a small RNA or not). As shown in Figure S4A, in the two-step RT-PCR analysis, theoretically, only the PCR amplification using P-2 as the initial primer will give the desired PCR product (Case II). As expected, a 62-nt PCR band was detected from the total RNA of the wild-type *C. acetobutylicum* when using the primer P-2 to initiate the PCR reaction (Figure S4B), indicating that the native transcriptional direction of the P*_2384_*_-140_ sequence. On this basis, the RACE experiment was carried out. The result further revealed a 94-nt transcript, which is located between the CAC2383 and CAC2384 genes and partially overlaps with the ORF of CAC2383 (Figure 3A). Given that this 94-nt short transcript has a stable and typical secondary structure (Figure 3B), no Shine-Dalgarno (SD) sequence, and start and stop codons, it was very likely as sncRNA. On this basis, Northern blotting using a single-stranded oligonucleotide probe targeting this 94-nt transcript was performed to further confirm the existence of this sncRNA. The whole DNA fragment covering the ORF of CAC2383 and CAC2384 as well as their intergenic region was PCR-amplified and then integrated it into a replicative plasmid for expression (Figure 3C), aiming to enrich the *in vivo* level of the potential sRNA. Encouragingly, after the resulting plasmid (psRNA) and a control plasmid (pControl) were transferred into *C. acetobutylicum* for Northern blot analysis, a desired approximate 94-nt hybridization signal was detected from the strain containing the plasmid psRNA, while no signal found from the control (Figure 3C).

**FIG 3.**
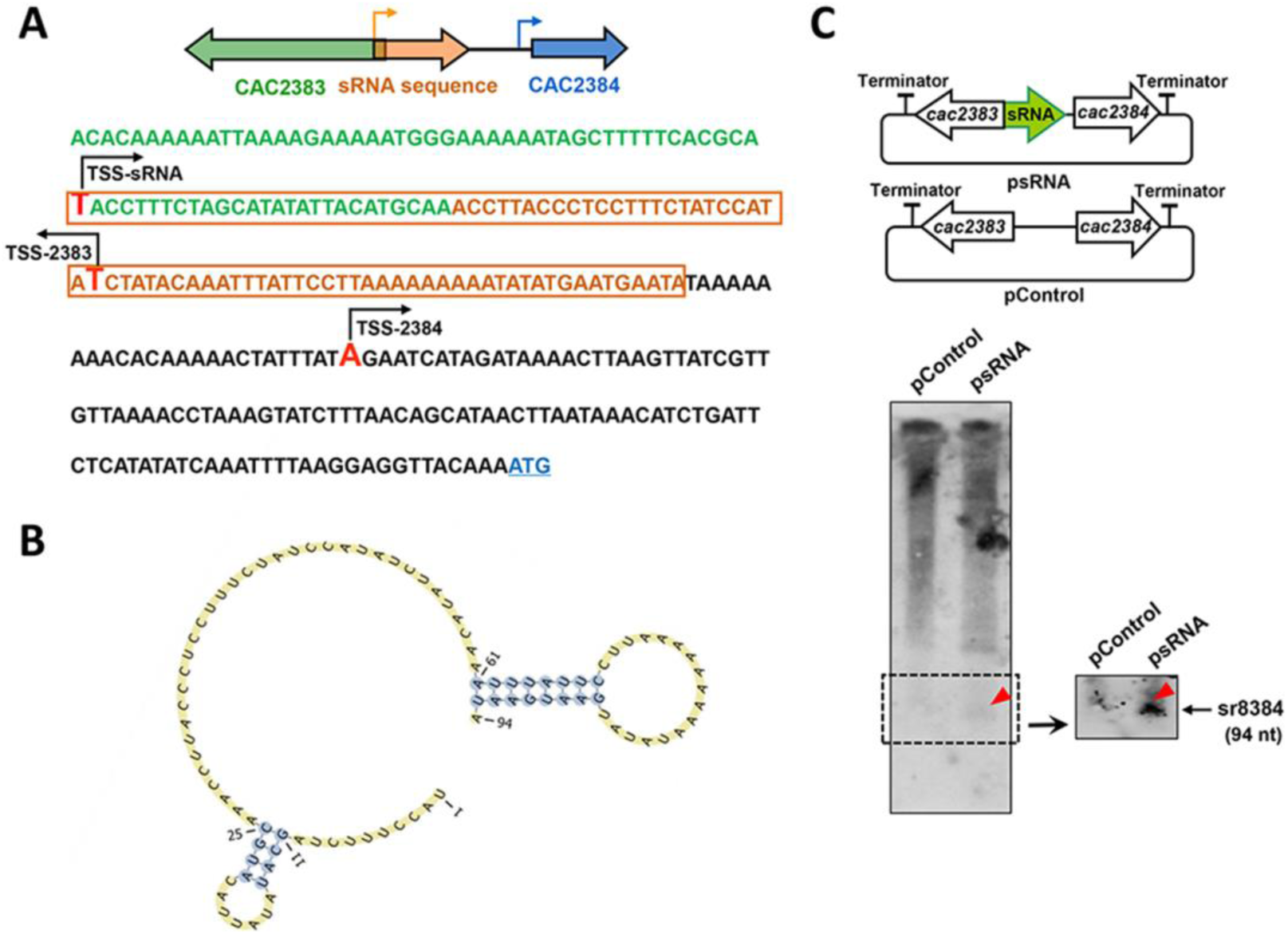
Sequence and structure of sr8384. (A) DNA sequence of CAC2383, CAC2384 and their intergenic region. The sr8384 sequence determined by the RACE reactions is shown in orange box. (B) The secondary structure of sr8384. (C) Northern blot analysis to identify the predicted sr8384. In this construct, the potential sncRNA-coding sequence was expressed by its native promoter; moreover, two terminators were located at two ends to avoid the potential expression running through from other genes on the plasmid.

In summary, the above results suggest the presence of a 94-nt sncRNA-coding sequence in the intergenic region between CAC2383 and CAC2384. Notably, this sncRNA, named sr8384 here, is not present in the list of sRNAs that were previously identified in *Clostridium* organisms *via* computational analysis (13), indicating that sr8384 has some novel genetic features.

### sr8384 is crucial for the control of cell growth and solvents synthesis in *C. acetobutylicum*

Having discovered the sr8384, it remained unknown whether this sncRNA is a crucial molecule in *C. acetobutylicum*. Therefore, the sr8384 transcript was disrupted for phenotypic examination using small regulatory RNA-based gene knockdown technology (18, 19). As shown in Figure 4A, a vector containing a 24-nt target-binding (TB) sequence that targets the middle region of sr8384 was constructed and introduced into the wild-type *C. acetobutylicum* strain, yielding the mutant strain 824(8384r). The mutant strain 824(8384r) exhibited a greater than 50% decrease in sr8384 transcript levels compared to the levels in the 824c strain (Figure 4C), demonstrating effective *in vivo* knockdown of the sr8384 transcript. Subsequently, in a batch fermentation, compared to the control strain 824c (containing the same plasmid lacking the 24-nt target-binding sequence), the 824(8384r) strain exhibited greatly impaired growth and synthesis of all the three major solvents (acetone, butanol and ethanol) (Figure 4B), in which the impact on butanol is especially significant (10.59 g/L vs. 14.54 g/L). These data suggest that sr8384 plays a crucial role in *C. acetobutylicum*.

**FIG 4.**
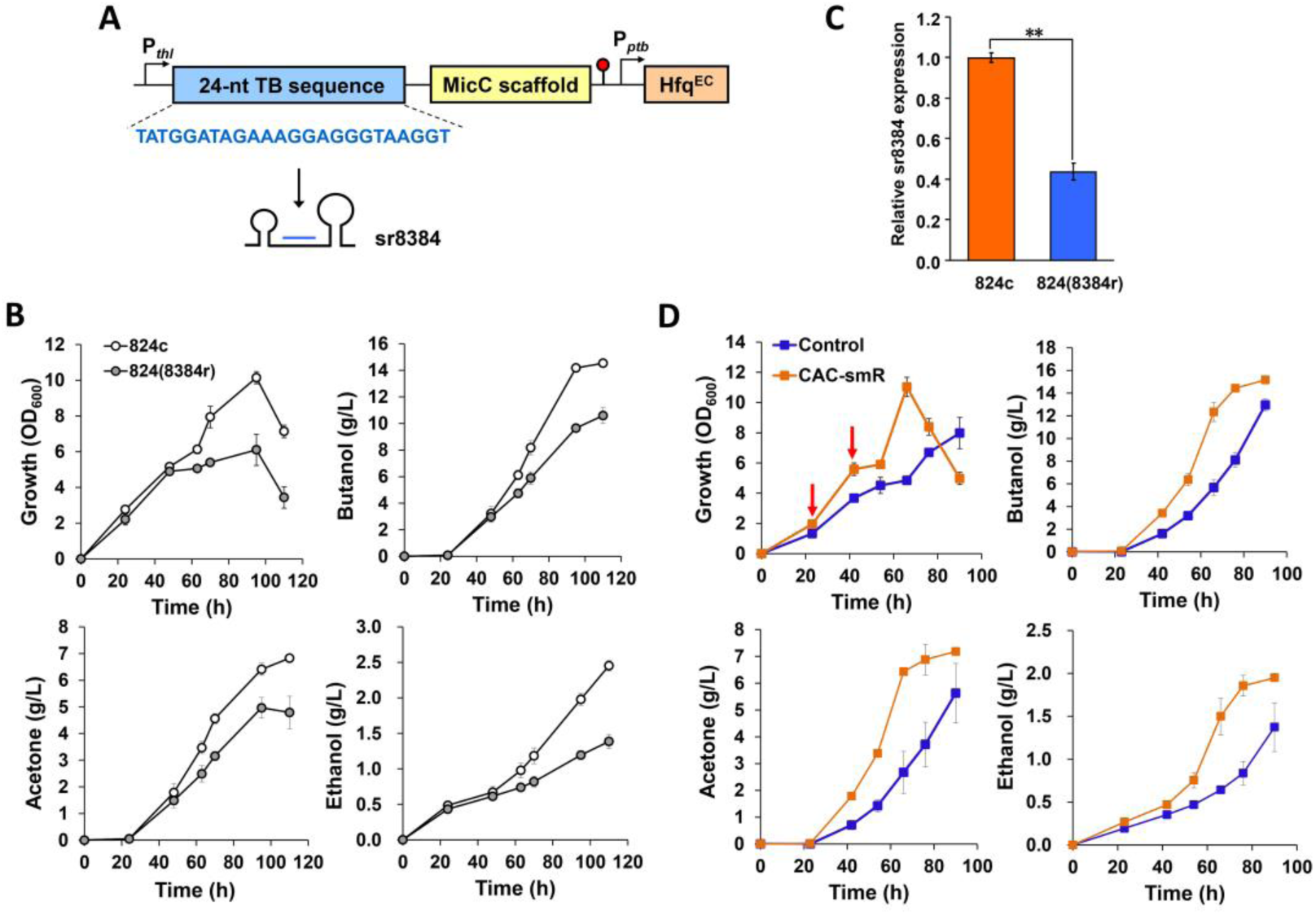
The influence of *in vivo* sr8384 level on cellular performance. (A) The construct of small regulatory RNA-based gene knockdown. TB: target-binding. The 24-nt TB sequence is responsible for targeting against sr8384. (B) Phenotypic effects of repressed sr8384 expression. 824c, 824c, the wild-type strain that carries the pIMP1-AS-con plasmid (without the 24-nt TB sequence); 824(8384r), the strain that carries the pIMP1-AS-sr8384 plasmid. Data are means±standard deviations of three independent experiments. (C) Fold change of sr8384 level in *C. acetobutylicum* after introducing antisense construct. (D) Phenotypic effects derived from increased sr8384 level. A 500 mL-working volume is used to perform the fermentation. The red arrows reflect the sampling time points (23 h and 42 h) for microarray assays. Control, the control strain that carries the empty plasmid; CAC-smR, the strain that carries the sr8384-overexpression plasmid pIMP1-P*_thl_*-sr8384. Data are means±standard deviations of two independent experiments.

Since sr8384 plays an essential role in *C. acetobutylicum*, a derived question is whether enhancement of the *in vivo* levels of this sncRNA could promote the cellular performance of *C. acetobutylicum*. Therefore, we constructed an expression vector in which the coding sequence of sr8384 was overexpressed under the control of a strong constitutive promoter, namely, P*_thl_*. The plasmid was then introduced into the wild-type *C. acetobutylicum*, yielding the strain CAC-smR. Encouragingly, the strain CAC-smR exhibited a greatly enhanced growth rate, biomass and production of the total solvents (acetone, butanol and ethanol) compared to the control strain (Figure 4D). Overall, these findings showcase not only the indispensability of sr8384 but also the potential value of this sncRNA as a molecular tool in *C. acetobutylicum*.

### Global regulatory role of sr8384 in *C. acetobutylicum*

Because sr8384 overexpression led to the positive phenotypic changes of *C. acetobutylicum* (Figure 4D), we used a comparative transcriptomics approach to search for genes affected by sr8384. The RNA samples for microarray assays were isolated from the sr8384-overexpressing strain CAC-smR and the control strain at two time points, namely, 23 h and 42 h, reflecting acidogenic and solventogenic stages, respectively (Figure 4D). The results showed that 679 and 380 genes exhibited significantly altered transcriptional levels (fold change ≥ 2.0) at 23 h and 42 h (Table S1 and S2), respectively, of which 172 genes were detected at both time points (Figure S5A). These differentially expressed genes could be roughly grouped into 16 subsets (Figure S5B), including some subsets of genes associated with important physiological and metabolic processes. These results indicate a crucial and global regulatory role of sr8384 in *C. acetobutylicum*. We selected 10 genes that exhibited different degrees of transcriptional repression or activation in the microarray assay after sr8384 overexpression for expression level validation using qRT-PCR. The qRT-PCR results were consistent with the data from the microarray analysis (Figure S6), indicating that the microarray data was of high quality.

Of note, five genes known to significantly influence the production of solvents in *C. acetobutylicum*, including a AbrB-coding gene, two histidine kinase-coding genes and two essential genes in the *sol* operon (4) were found to be significantly upregulated after sr8384 overexpression according to the microarray data (Figure S7A), although the results of RNA hybridization analysis showed no binding activity between sr8384 and the transcripts of these five genes (Figure S7B). Therefore, it can be concluded that sr8384 indirectly activates the expression of these crucial genes, which may contribute to the solvent production in *C. acetobutylicum*.

Our next challenge was to identify direct targets controlled by sr8384. To this end, we first used the online tool IntaRNA (20) to predict putative target sequences based on their potential interaction energy with sr8384. The top 100 sequences (with interaction energies ≤ −18.3083 kcal/mol) within the predicted results were chosen for further investigation. The genes associated with these 100 sequences (located in the promoter or coding region) that exhibited ≥ 2-fold transcriptional changes (microarray assay) after sr8384 overexpression were selected, resulting in 26 candidates as well as their associated genes being used for further detailed investigations (Figure 5A). As shown in Table S3, most of these 26 target sequences spanned both the promoter and coding regions of their corresponding genes. Next, these 26 candidates were used for RNA hybridization analysis to examine whether these genes interact with sr8384. The results showed that, of the 26 candidates, 15 exhibited distinct binding activity with sr8384 (Figure 5B), whereas no obvious binding was observed for the remaining candidates. Among these 15 sequences of direct targets of sr8384, nine were associated with genes with annotated functions (Table S3).

**FIG 5.**
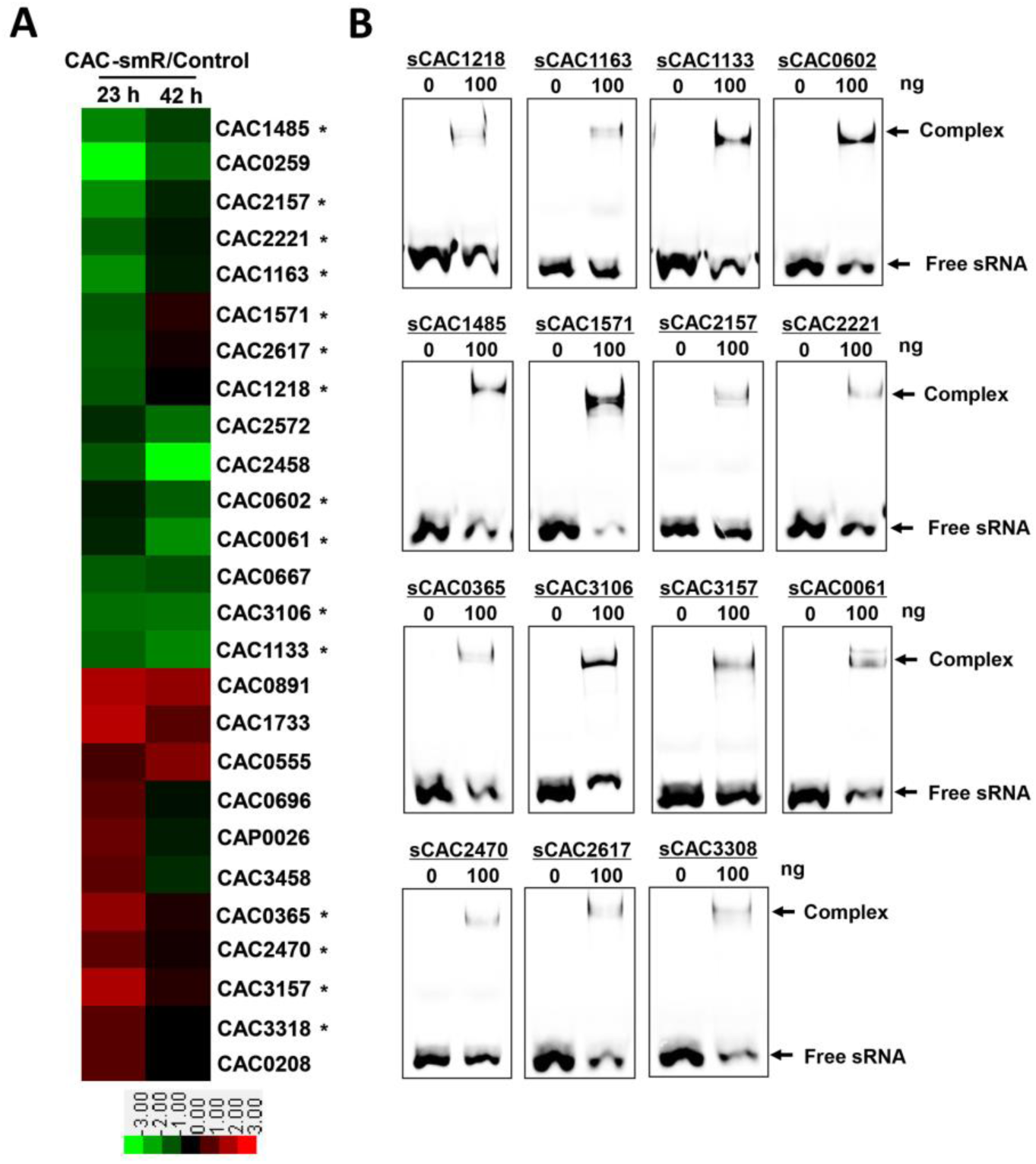
Identification of the direct targets of sr8384 in *C. acetobutylicum*. (A) The 26 picked genes that are potentially controlled by sr8384 and simultaneously showed over 2-fold expressional changes after sr8384 overexpression. (B) 15 identified target sequences directly bind with sr8384. These 15 genes directly regulated by sr8384 were labelled with asterisk in (A).

To explore whether the genes associated with these 15 sr8384 target sequences contributed to the phenotypic changes of the CAC-smR strain, we conducted knockdown or overexpression of these genes (11 knockdown strains and 4 overexpressing strains) (Figure 6), according to the transcriptional alterations of these genes after sr8384 overexpression (Figure 5A), yielding a total of 15 mutant strains.

**FIG 6.**
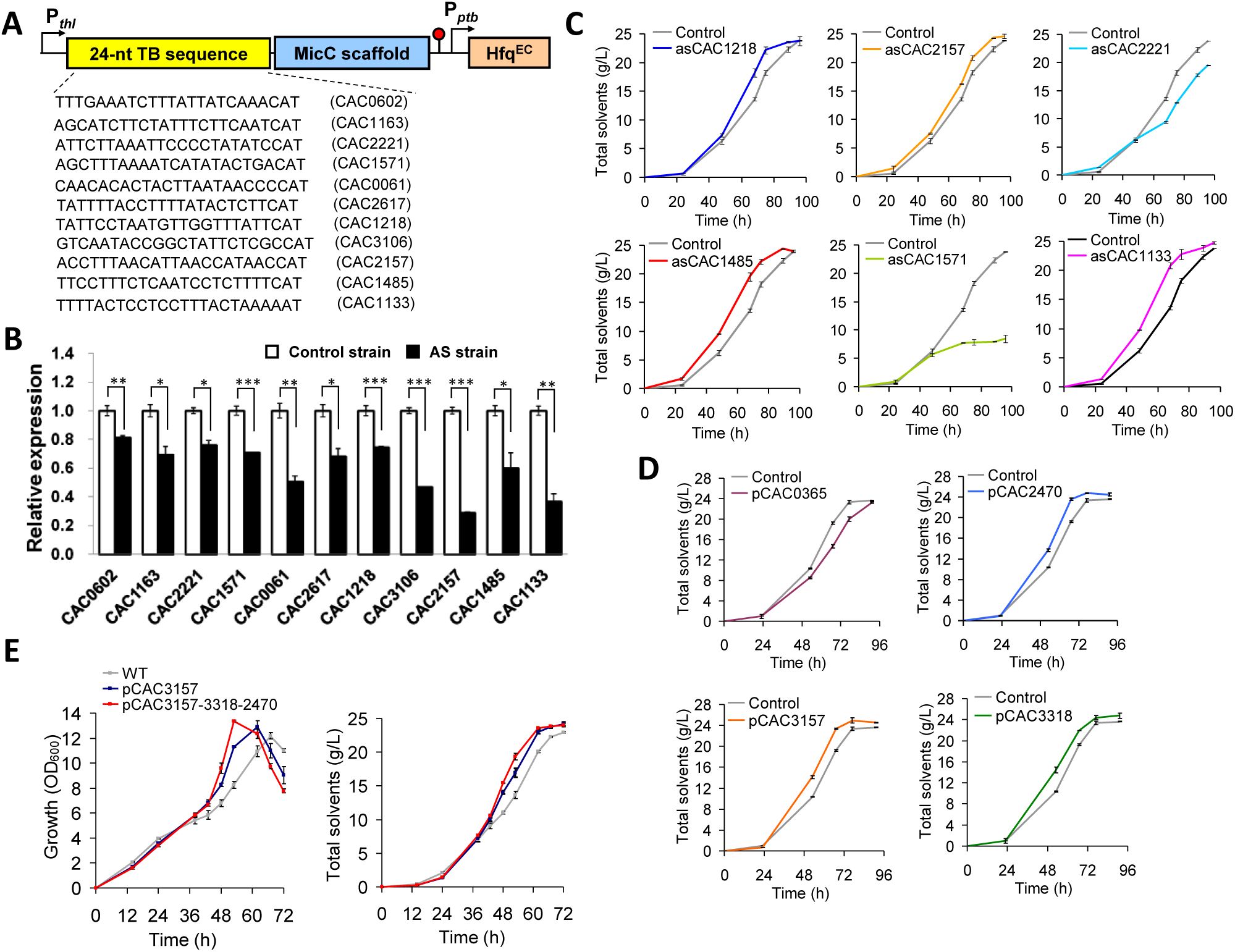
Functional roles of the 15 direct sr8384 targets in *C. acetobutylicum*. (A) The constructs for small regulatory RNA-based knockdown of the 11 target genes of sr8384. TB: target-binding. The *E. coli hfq* gene was expressed under the control of *ptb* promoter (B) The 11 sr8384 targets with repressed expressional levels by antisense RNA. (C) Altered solvent production derived from expressional repression of CAC1218, CAC2157, CAC2221, CAC1485, CAC1571 and CAC1133. (D) Altered solvent production derived from the overexpression of CAC2470 (pCAC2470), CAC3157 (pCAC3157), CAC3318 (pCAC3318) and CAC0365 (pCAC0365). (E) Comparison of the solvents production between single CAC3157 overexpression (pCAC3157) and combined overexpression of CAC3157, CAC3318 and CAC2470 (pCAC3157-3318-2470). Control, the *C. acetobutylicum* strain harboring a blank plasmid skeleton. Data are means±standard deviations of three independent experiments.

Here, the knockdown of the 11 genes was performed by using the same method mentioned above (18, 19), in which a 24-nt sequence targeting each gene was expressed (Figure 6A). By this method, the transcriptional levels of the 11 genes were decreased to different extents (Figure 6B), and the resulting mutants were used for phenotypic examination. Only 6 of the 11 genes, namely, CAC1218, CAC2157, CAC2221, CAC1485, CAC1571 and CAC1133, caused increased or decreased synthesis of the solvents after their knockdown (Figure 6C and S8A). Notably, knockdown of CAC1571 significantly impaired solvent production (Figure 6C). CAC1571 is annotated to encode glutathione peroxidase, an enzyme known to protect organisms from oxidative damage (21). Thus, repression of the expression of this gene may damage the basic tolerance of *C. acetobutylicum* to oxygen stress during anaerobic fermentation.

Upon overexpression of the other four genes, namely, CAC2470, CAC3157, CAC3318 and CAC0365, increased or accelerated solvents formation was observed for the first three strains (pCAC2470, pCAC3157 and pCAC3318), while a clear lag in solvent synthesis was observed for the final strain (pCAC0365) (Figure 6D and S8B). The CAC0365 gene is predicted to encode a phosphoglycerate dehydrogenase, an enzyme that catalyzes the synthesis of serine from 3-phosphoglycerate (22). Thus, as shown in Figure S9, overexpression of CAC0365 could decrease the metabolic flux from 3-phosphoglycerate to pyruvate, the precursor for solvent synthesis, thereby impairing solvent production in *C. acetobutylicum*.

Because separate overexpression of CAC2470, CAC3157 or CAC3318 all led to increased solvent production in *C. acetobutylicum*, we asked whether the combined overexpression of these functional genes would have a synergistic effect. To this end, we coexpressed these three genes under the control of the constitutive promoter P*_thl_*. As shown in Figure 6E and S8C, the engineered strain pCAC3157-3318-2470 further exhibited slightly increased growth and solvent production compared to the strain pCAC3157 (the one with highest solvent titer among the three separate overexpressing strains as shown in Figure 6D). This finding confirms the above hypothesis and, moreover, indicates that a combined modulation of the sr8384 targets may further improve the cellular performance of *C. acetobutylicum*..

### CAC2385: an indirect sr8384 target with a significant effect on cell growth and solvent production

According to the microarray data, we observed that the CAC2385 gene, which is located downstream of the sr8384 sequence (453 nt apart) in the chromosome, showed a nearly 3-fold upregulation (at 23 h) after sr8384 overexpression (Table S1), suggesting a potential regulatory effect of sr8384 on CAC2385. However, RNA hybridization analysis showed no direct interaction between sr8384 and the transcript of CAC2385 or its upstream noncoding region (Figure 7A). Therefore, the altered CAC2385 expression after sr8384 overexpression (Table S1) was likely due to an indirect effect.

**FIG 7.**
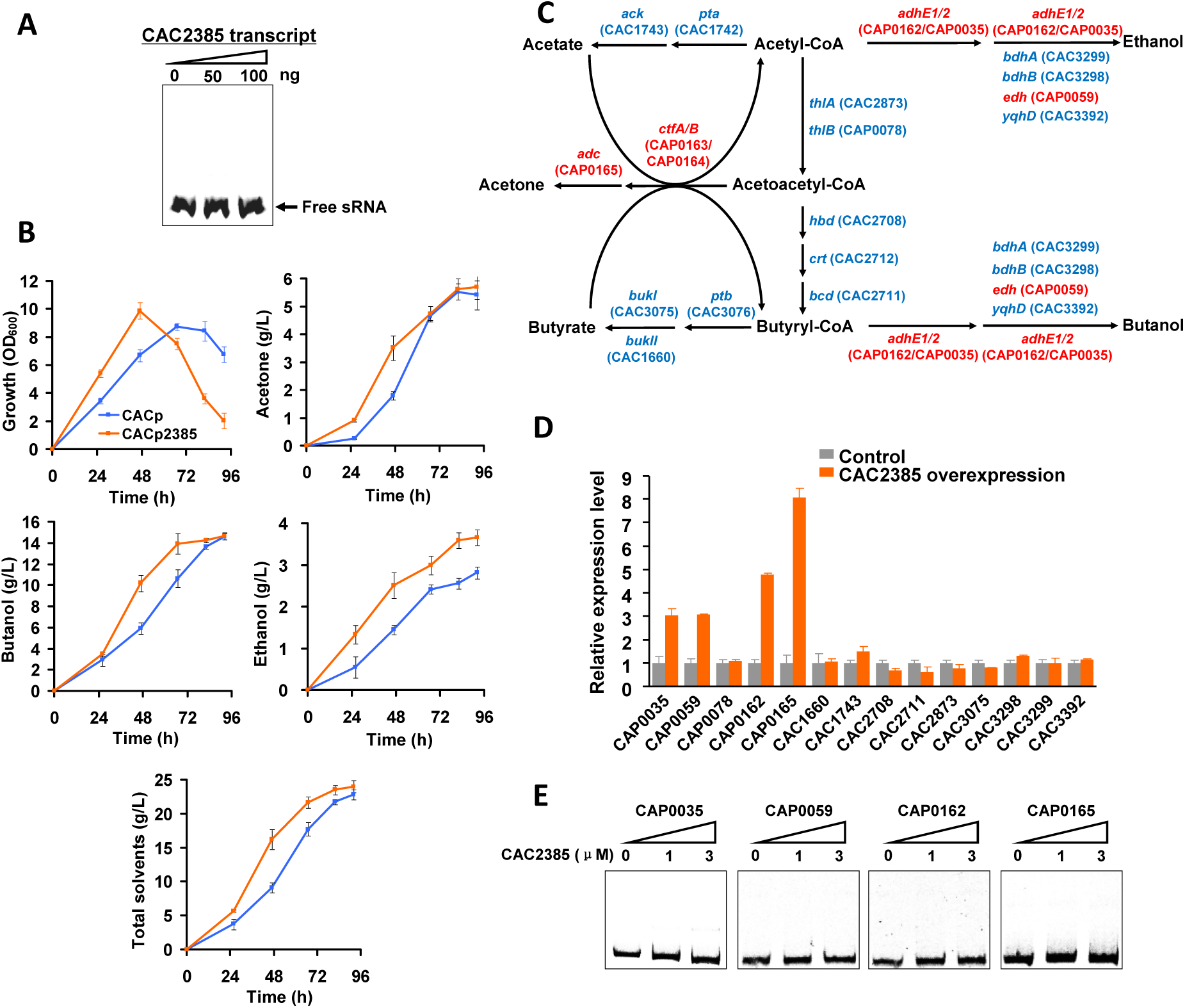
Characterization and functional analysis of the CAC2385 gene. (A) RNA hybridization analysis of sr8384 and the CAC2385 transcript (covering the coding and the upstream noncoding region of CAC2385). (B) Phenotypic changes derived from CAC2385 overexpression. CACp, the strain carries the pIMP1-P*_ptb_* plasmid; CACp2385, the strain carries the pIMP1-P*_ptb_*-CAC2385 plasmid. Data are means±standard deviations of three independent experiments. (C) All essential genes located in solvents synthetic pathways in *C. acetobutylicum*. (D) The expressional changes of the genes that located in solvents synthetic pathways after CAC2385 overexpression. The genes activated by CAC2385 were labelled as red in (C). Here, we used the expressional level of CAP0162 to represent that of the *sol* operon (cotranscribed CAP0162-0163-0164). Data are means±standard deviations of two independent experiments. (E) EMSAs of binding of CAC2385-coding protein to the promoter region of CAP0162, CAP0035, CAP0059 and CAP0165.

However, the function of the CAC2385 gene is not known. When SMART (23) (Simple Modular Architecture Research Tool, http://smart.embl-heidelberg.de) was used to analyze the protein encoded by CAC2385, a DNA-binding HTH domain was predicted, indicating that this protein is likely a transcription factor (TF). To explore the role of CAC2385 in *C. acetobutylicum*, this gene was overexpressed for phenotypic investigation. Encouragingly, the resulting strain (CACp2385) exhibited significantly improved cellular performance compared to the performance of the control strain (CACp), i.e., a much higher growth rate and increased productivity of the three major products (Figure 7B). This finding suggests that the CAC2385 gene constitutes an important element in the sr8384 regulatory network.

Based on the significantly enhanced solvent production after CAC2385 overexpression, we specifically examined the transcriptional changes associated with all the essential genes in solvent synthetic pathways (Figure 7C) using qRT-PCR. The results showed that the *adhE1* gene (CAP0162); two major alcohol dehydrogenase-coding genes, namely, *adhE2* (CAP0035) and *edh* (CAP0059); and the acetone synthetic pathway gene *adc* (CAP0165) were all greatly upregulated (4.78, 3.03, 3.08 and 8.08-fold increase, respectively) (Figure 7D). Actually, the expression level of *adhE1* can represent that of the *sol* operon, which is a cotranscribed gene cluster (CAP0162-0163-0164). The greatly enhanced expression of these essential genes responsible for solventogenesis may partly explain the significantly increased ability of the strain CACp2385 in the production of acetone, ethanol and butanol. However, no binding activity was detected between the CAC2385 protein and intergenic regions upstream of the abovementioned genes (CAP0162, CAP0035, CAP0059 and CAP0165) according to the EMSA results (Figure 7E), indicating that the influence of CAC2385 to these targets is also indirect.

### Manipulation of sr8384 and its homolog leads to improved growth and solvent synthesis in Clostridium

To further explore the role of sr8384 in genetic improvement based on the above result (Figure 4D), we used three promoters (P_200-1_, P*_thl_* and P_1200-9-15_) (24) with gradually increasing activities for sr8384 overexpression (Figure 8A) to determine whether this sncRNA has a dosage-dependent effect on cellular performance. As expected, the resulting three strains exhibited improved growth rate and solvent titer and productivity to different extents compared to the control strain (Figure 8B); however, surprisingly, the best effect was seen with the weakest promoter, namely, P_200-1_, rather than the other two stronger promoters (Figure 8B), reflecting that the *in vivo* sr8384 level is associated with the cellular performance of *C. acetobutylicum*.

**FIG 8.**
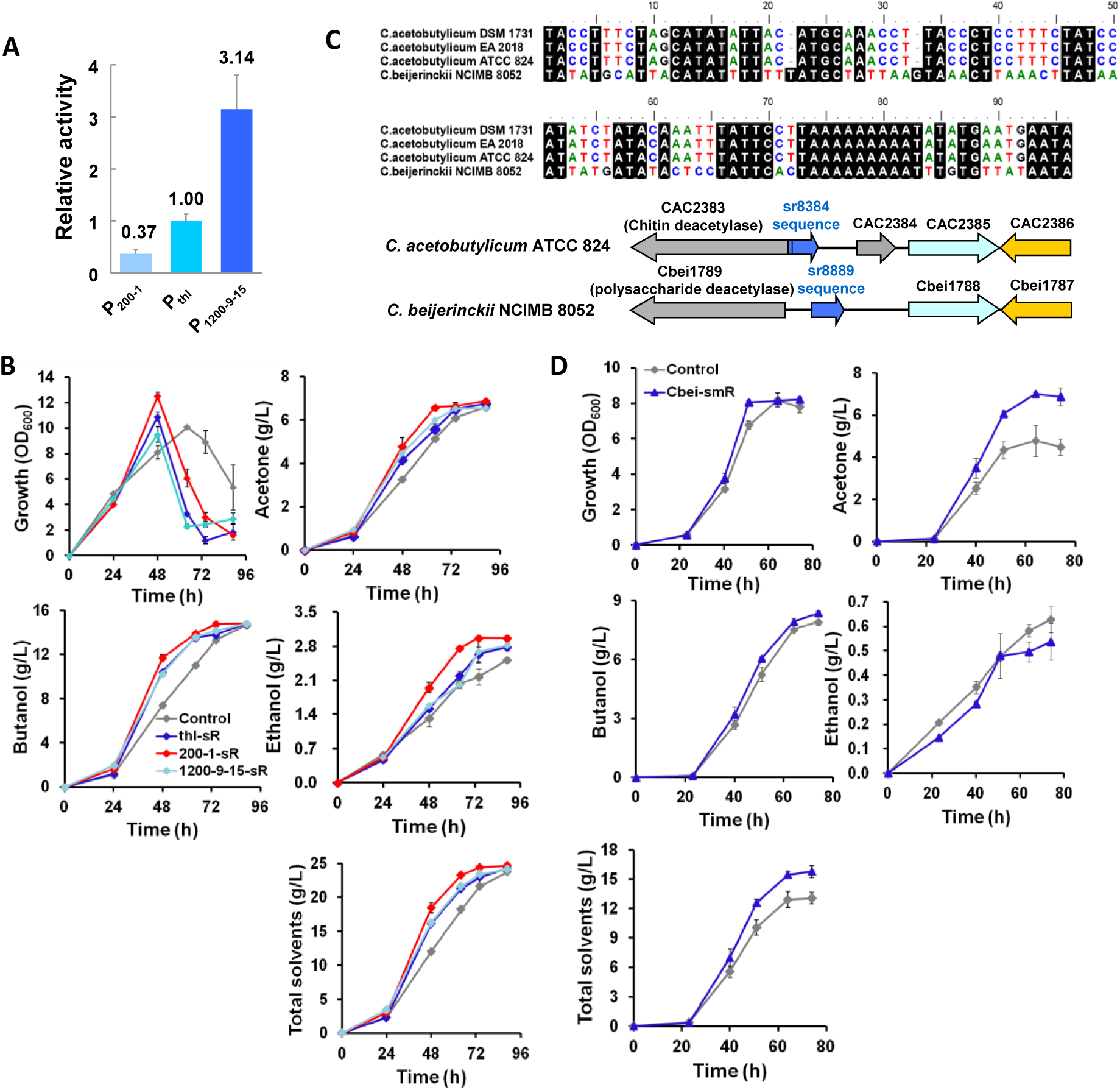
Manipulation of sr8384 and its homolog can improve the growth and solvent synthesis of *Clostridium*. (A) Three promoters with gradually increased activities used for sr8384 overexpression. (B) Improved growth and total solvents by sr8384 overexpression using the three promoters in (A). Control, the control strain harbours the empty plasmid; thl-sR, the strain with sr8384 overexpressed under the promoter of P_thl_; 200-1-sR, the strain with sr8384 overexpressed under the promoter of P_200-1_; 1200-9-15-sR, the strain with sr8384 overexpressed under the promoter of P_1200-9-15_. (C) The coding sequence of sr8384 homologs in clostridia. (D) Improved cellular performance of *C. beijerinckii* by sr8889 overexpression (the sr8384 homolog in *C. beijerinckii*). Data are means±standard deviations of three independent experiments.

To date, numerous sRNAs have been found to be conserved in several genera (25). Thus, we sought to find sr8384 homologs in other clostridial genome sequences available in NCBI. BlastN analysis showed that no putative sr8384 homologs were present in any other *Clostridium* species except two *C. acetobutylicum* strains (EA2018 and DSM1731) (Figure 8C). Interestingly, when we scanned the genome of *C. beijerinckii* NCIMB 8052, another major solventogenic *Clostridium* species, a potential homologous sequence that shares high identity (54.1%) with the sr8384 sequence was found in the intergenic region between the Cbei1789 and Cbei1788 genes (which are two conserved orthologs in *C. beijerinckii*, corresponding to CAC2383 and CAC2385 in *C. acetobutylicum*, respectively) (Figure 8C). Here, this sequence was named sr8889. Next, we investigated whether sr8889 was also functional in *C. beijerinckii*. As expected, overexpression of this sequence indeed resulted in an increased growth rate and a greater than 20% increased titer of the total solvents (acetone, butanol and ethanol) compared with the values for these parameters in the wild-type strain (Figure 8D). Taken together, these results show that sr8889 plays an important role in *C. beijerinckii*. Moreover, although not highly conserved in different *Clostridium* species, this type of pleiotropic sncRNA appears to be crucial for industrial clostridia, which has not been fully recognized and remains to be explored.

## DISCUSSION

Discovery and functional analysis of sncRNAs have been performed in some representative industrial microorganisms, such as *E. coli* (26), *Saccharomyces cerevisiae* (27, 28) and *B. subtilis* (5, 29). However, sncRNAs as well as their potential values in metabolic engineering remain largely unexplored in solventogenic *Clostridium* species. In this study, we discovered the atypical sncRNA sr8384 and its utility in the control of growth and solvent synthesis in *C. acetobutylicum*, a model organism for clostridia. To the best of our knowledge, such a determinant sncRNA with a crucial regulatory role in clostridia has not been previously reported.

To date, only an extremely limited number of sncRNAs have been shown to be associated with certain functions in clostridia (10–12). The sncRNA sr8384 identified here is a global rather than specific regulatory molecule in *C. acetobutylicum*. Given that sr8384 was not predicted or detected in the previous screenings for sRNAs in clostridia based on comparative genomics and RNA-seq (13, 30, 31), this sncRNA is likely atypical in sequence or is very poorly expressed in *C. acetobutylicum*. Moreover, sr8384 exhibited a dose-dependent effect in the regulation of phenotypes of *C. acetobutylicum*, i.e., negative and positive phenotypic changes were observed upon repression and overexpression, respectively, of this sncRNA (Figure 4B and D), thus indicating that sr8384 has potential application as a target for strain improvement.

The greatly enhanced growth rate and solvent productivity upon sr8384 overexpression are two important phenotypic alterations in *C. acetobutylicum*. This effect could be attributed to the direct or indirect regulation by sr8384 of multiple effective targets (Figure 6C and D). Among the 10 sr8384 targets involved in the effect on solvent production (Figure 6C and D), CAC1571 exhibited the most negative changes when repressed (Figure 6C). As mentioned above, CAC1571 encodes a putative glutathione peroxidase. This enzyme is essential in organisms and is capable of protecting cells from oxidative damage (21, 32). In *C. acetobutylicum*, in addition to CAC1571, two other genes (CAC1570 and CAC1549) were also predicted to encode glutathione peroxidase (33). All three of these genes have been found to be rapidly upregulated in response to O_2_ flushing (21), indicating the importance of these genes in the scavenging of reactive oxygen species (ROS) in the anaerobic *C. acetobutylicum*. Therefore, the phenotypic changes caused by the repression of CAC1571 further reinforce the above hypothesis.

Notably, among the 11 sr8384 targets that were chosen for phenotypic investigation using small regulatory RNA interference (Figure 6A), some were not greatly knocked down at the transcript level (Figure 6B). For example, the transcriptional levels of CAC0602, CAC1163 and CAC2617 were downregulated by only less than 30%. The insufficient repression of these genes may not truly reveal their effects on cellular phenotypes, and detailed investigations remain to be performed.

As mentioned above, the *sol* operon is crucial for acid (acetic and butyric acids) assimilation and ABE solvent formation in *C. acetobutylicum*. Significant upregulation (6.57-fold) of the *sol* genes was also observed after sr8384 overexpression (Table S1). Therefore, upregulation of the *sol* genes definitely contributes to the enhancement in solvent production derived from sr8384 overexpression.

In addition, we observed that the transcriptional levels of CAC3319 and CAC0323, two orphan histidine kinase-coding genes, increased 2.16- and 2.24-fold, respectively, after sr8384 overexpression (Table S1). These two genes are known to be responsible for the phosphorylation of Spo0A. This protein is a well-known global regulator involved in the control of multiple physiological and metabolic processes and is simultaneously capable of improving solvent production by activating several key genes (the *sol* operon, *adc*, *bdhA* and *bdhB*) in *C. acetobutylicum* (34, 35). Therefore, we believe that regulation of the CAC3319 and CAC0323 genes by sr8384 constitutes an important part of the entire regulatory network of this sncRNA.

It should be noted that, in addition to the above-identified targets, sr8384 might regulate other genes. Here, although the comparative transcriptomic data for the sr8384-overexpressing strain in combination with the application of the online tool IntaRNA has proven useful for the prediction of sncRNA targets in *C. acetobutylicum*, it cannot be ruled out that some additional targets may have been missed. For example, the expression of some targets might change only when sr8384 is absent, rather than overexpressed, or some targets might not be expressed in the presence of D-glucose due to the CCR (carbon catabolite repression) effect. Therefore, for a comprehensive understanding of the global regulatory function of sr8384, microarray analyses based on sr8384 deletion or using other major carbon sources are necessary.

In summary, our data here identify sr8384 as a pleiotropic regulator in *C. acetobutylicum*. Multiple direct and indirect targets that are associated with different essential biological processes in *C. acetobutylicum* are controlled by this sncRNA, revealing a preliminary regulatory network (Figure 9). This pleiotropic function enables sr8384 to play a determinant role in *C. acetobutylicum*. Notably, improved cellular performance was achieved via modulation of the expression of sr8384 or its target genes. With these characteristics, sr8384, to the best of our knowledge, is the first identified sncRNA involved in regulating various physiological and metabolic processes in the industrially important *Clostridium* species. In addition, given the discovery of a functional sr8384 homolog in *C. beijerinckii*, another important industrial *Clostridium* species, this type of functional sncRNA may be widespread in clostridia, including pathogenic *Clostridium* species. In summary, this work provides new insight into the role of sncRNAs in clostridia and offers new opportunities for engineering these anaerobes.

**FIG 9.**
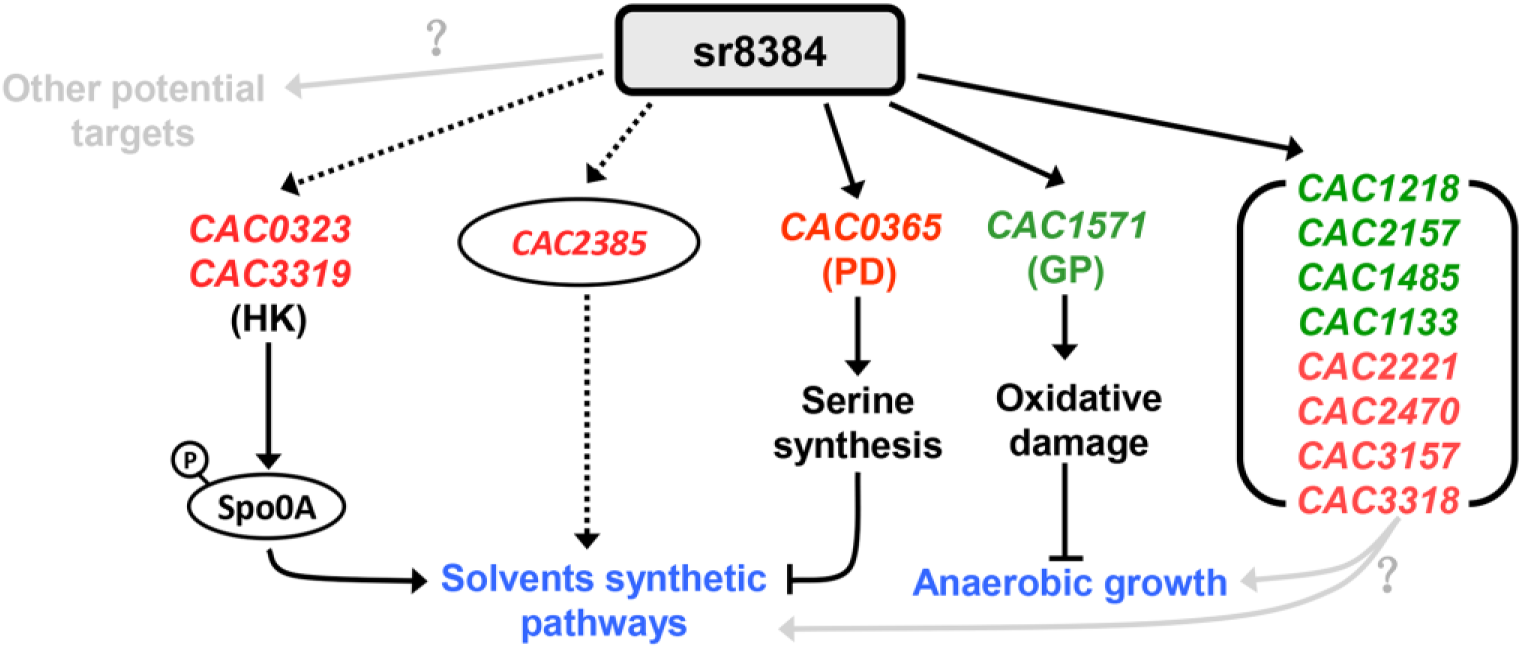
Schema illustrating the pleiotropic regulation of sr8384 in *C. acetobutylicum*. The dotted and solid arrow mean indirect and direct regulation, respectively. Spo0A: the global regulator that is involved in controlling spore formation and many other physiological and metabolic processes in clostridia. Both CAC0323 and CAC3319 encode histidine kinase responsible for Spo0A phosphorylation. PD: phosphoglycerate dehydrogenase. GP: glutathione peroxidase. The genes shown in red and green represent expressional activation and repression by sr8384, respectively. The questions remained to be explored were marked with interrogation points.

## MATERIAL AND METHODS

### Media and cultivation conditions

Luria-Bertani (LB) medium, supplemented with ampicillin (100 µg/mL) and spectinomycin (50 µg/mL) when needed, was used to cultivate *E. coli*. *C. acetobutylicum* was first grown anaerobically (Thermo Forma Inc., Waltham, MA) in CGM medium (36) for inoculum preparation. Upon reaching the exponential growth phase (OD_600_=0.8-1.0), the cells (5% inoculation amount) were transferred into P2 medium (37) for fermentation. Erythromycin (10 µg/mL) and thiamphenicol (8 µg/mL) were added to the P2 medium when needed. Samples for assays were removed at different time points and then stored at −20°C.

### Bacterial strains and plasmid construction

The primers used in this study are listed in Table S4. The strains and plasmids used in this work are listed in Table S5. Top10 cells were used for gene cloning. The plasmids were first methylated by *E. coli* ER2275 and then electroporated into *C. acetobutylicum*.

The CAC2384 gene was disrupted using the group II intron-based Targetron system. In brief, a 350-bp DNA fragment was amplified by PCR with the following primers: the EBS universal primer, CAC2384-174,175s-IBS, CAC2384-174,175s-EBS1d and CAC2384-174,175s-EBS2. Amplification was performed according to the protocol of the Targetron Gene Knockout System Kit (Sigma-Aldrich, St Louis, MO, USA). After digestion with *Xho*I and *BsrG*I, this 350-bp DNA fragment was cloned into the plasmid pWJ1 (38), yielding the plasmid pWJ1-CAC2384.

The pP*_thl_* plasmid was constructed as previously reported (39). The P*_2384_* fragment was amplified by PCR using the primers P*_2384_*-for and P*_2384_*-rev with the genomic DNA of *C. acetobutylicum* as the template. After digestion with *Pst*I and *BamH*I, the P*_2384_* fragment was cloned into the plasmid pP*_thl_*, yielding the plasmid pP*_2384_*. Similarly, the CAC2384 fragment was amplified by PCR using the primers CAC2384-for and CAC2384-rev. Then, the CAC2384 fragment was digested with *Sal*I and *BamH*I and ligated to the plasmids pP*_thl_* and pP*_2384_*, yielding the plasmids pP*_thl_*-2384 and pP*_2384_*-2384, respectively. The plasmid pIMP1-f62 was derived from pP*_thl_* by replacing the promoter P*_thl_* with P*_2384-140_*. The pIMP1-f62-LT and pIMP1-f62-RT plasmids were derived from pIMP1-f62 by adding the terminator at the left side or right side of P*_2384-140_*, respectively. The plasmid pIMP1-P*_ptb_*-CAC2385 was generated from pIMP1-P*_ptb_* (40) by adding the CAC2385 gene under the control of the promoter P*_ptb_*.

For overexpression of sr8384 and sr8889 in *C. acetobutylicum* and *C. beijerinckii*, respectively, the sr8384 and sr8889 sequences were amplified by PCR, digested with *Sal*I and *BamH*I, and then ligated with the plasmid pP*_thl_* that had been digested with the same restriction enzymes, yielding the plasmids pIMP1-P*_thl_*-sr8384 and pIMP1-P*_thl_*-sr8889, respectively.

The plasmid used for small regulatory RNA-based knockdown of sr8384 was constructed according to a previously reported method (18, 19). In brief, the fragment P*_thl_*-AS-sr8384-MicC, which contained a 24-nt target-binding sequence complementary to sr8384 and the MicC sRNA scaffold, was first amplified by PCR using the plasmid pP*_thl_* as the template with the primers AS-P*_thl_*-s and AS-sr8384-MicC-a. Then, a fragment containing both the promoter P*_ptb_* and Hfq^EC^ (P*_ptb_*-Hfq^EC^) was obtained by overlap PCR. Finally, the fragments P*_thl_*-AS-sr8384-MicC and P*_ptb_*-Hfq^EC^ were assembled by overlap PCR, yielding the large fragment P*_thl_*-AS-sr8384-MicC-P*_ptb_*-Hfq^EC^. After digestion with *Pst*I and *EcoR*I, the fragment P*_thl_*-AS-sr8384-MicC-P*_ptb_*-Hfq^EC^ was ligated with the plasmid pP*_thl_* that had been digested with the same restriction enzymes, yielding the plasmid pIMP1-AS-sr8384. The method for the construction of the plasmids used for knockdown of the other target genes in this study was the same as that used for sr8384, changing only the 24-nt target-binding sequence.

For overexpression of CAC0365, CAC3157, CAC2470 and CAC3318, these genes were firstly PCR-amplified with the primers listed in Table S4. After digestion with *Sal*I and *BamH*I, these fragments were ligated to pP*_thl_* to yield the plasmids pP*_thl_*-CAC0365, pP*_thl_*-CAC3157, pP*_thl_*-CAC2470 and pP*_thl_*-CAC3318. Next, to study the synergistic effect of CAC3157, CAC2470 and CAC3318 on growth and solvents production of *C. acetobutylicum*, CAC3318 and CAC2470 were PCR-amplified separately with the primers listed in Table S4. Then, CAC3318-CAC2470 was acquired by overlap PCR with CAC3318 and CAC2470 supplied as templates. After digestion with *BamH*I and *EcoR*I, CAC3318-CAC2470 was cloned to the plasmid pP*_thl_*-CAC3157, yielding pP*_thl_*-CAC3157-3318-2470.

### Gene disruption and functional complementation using a group II intron **(targetron)**

Gene disruption in *C. acetobutylicum* was achieved through chromosomal insertion of a group II intron by using the targetron plasmid (15). The primers used for retargeting the RNA portion of the intron to target genes were listed in Table S4. In detail, the fragments of P*_thl_*, P*_thl_*-CAC2384, P*_2384_*, P*_2384_*-CAC2384 and the two separate part of the initial intron sequence were PCR-amplified by using the primer pairs listed in Table S4. Then, the whole functional sequence, which contained both the intron and the insertion sequence, were acquired by overlap PCR with Intron-for/Intron-rev supplied as primers and inserted into the chromosome by using the “targetron” strategy.

### Analytical methods

The density of the culture (A_600_) after cell growth was tested using a spectrophotometer (DU730, Beckman Coulter, Placentia, California, USA). The concentrations of the solvents (acetone, acetic acid, butyric acid, butanol, and ethanol) were determined using gas chromatography (7890A, Agilent, Wilmington, DE, USA). Isobutyl alcohol and isobutyric acid were used as the internal standards for solvent quantification.

### Identification of transposon insertions in the chromosome by reverse transcription PCR

The transposon mutant library of *C. acetobutylicum* that was constructed according to a previously reported protocol (14) was used to identify mutants that exhibited greatly changed solvent production. The selected mutant strain was used for reverse transcription PCR analysis to identify the transposon insertion site in the chromosome as reported previously (14).

### Two-step RT-PCR analysis

The two-step RT-PCR analysis was performed according to the previous report (41), in order to determine the transcriptional direction of the potential small RNA sr8384. In brief, the total RNA of *C. acetobutylicum* was isolated by TRIzol extraction. A pair of primers that match the sr8384 sequence were separately added into the total RNA to synthesize the first-strand cDNA by using the PrimeScript RT Reagent Kit (TaKaRa, cat. #RR047A). Next, two primers were simultaneously used for the second round of PCR amplification using the above first-strand cDNA as the template. Finally, the PCR products were separated on 1.5% agarose gels.

### Southern blot analysis

Southern blot was performed using a digoxigenin (DIG) High Prime DNA Labeling and Detection Starter Kit I (Roche Diagnostics GmbH, Mannheim, Germany) as instructed by the manufacturer. Briefly, 10 µg of genomic DNA was digested with *Hind*III for about 14 h, separated on a 1.0% agarose gel, and then transferred to a charged nylon membrane. Next, the digoxigenin-labeled DNA probe was used for southern hybridization.

### Northern blot analysis

In the Northern blot analysis, 50 μg of total RNA was loaded and electrophoretically resolved on a 7% denaturing polyacrylamide gel containing 7M urea. Then, the RNA was transferred to an Immobilon-NY+ membrane (Merck KGaA, Darmstadt, Germany, INYC00010) and immobilized by UV-crosslinking. A 94-nt digoxigenin-labeled probe (20 pmol) complementary to the sRNA sequence was synthesized and used to detect the presence of the sRNA. The prehybridization (1-2 h) and hybridization (16 h) steps were performed at 37°C using NorthernMax^®^ prehybridization and hybridization buffers (LifeTech, Thermo Fisher Scientific Inc., Carlsbad, CA, USA; cat: AM8677). The membrane was washed twice with 4×SSC buffer for 15 min and then washed with 2×SSC buffer (2×SSC, 0.1% SDS) for an additional 15 min. Finally, immunological detection was performed according to the protocol for the DIG Northern Starter Kit (Roche, Mannheim, Germany, Cat. No. 12039672910).

### 5’ and 3’ rapid amplification of cDNA ends (RACE) analysis

The 5’ and 3’ RACE analyses were performed according to the protocol for the SMARTer^®^ RACE 5’/3’ Kit (TaKaRa Bio USA, Inc., cat. nos. 634858, 634859). The whole upstream noncoding region (202 nt) of the CAC2384 gene, together with the ORF of CAC2383 and CAC2384, was PCR-amplified from the chromosome of *C. acetobutylicum* and then integrated into a multicopy plasmid to enrich the transcripts for RACE. For the 3’ RACE analysis, a poly(A) tail was first added to the 3’ end of the RNA template using *E. coli* poly(A) polymerase (New England BioLabs, Beverly, MA, USA; M0276S).

### Microarray analysis

CAC-smR and the control strain were grown in P2 medium (500 ml) with erythromycin supplementation (10 µg/mL). D-glucose (80 g/L) was used as the sole carbon source. Samples for microarray analysis were taken at 23 h (acidogenic phase) and 42 h (acid-solvent transition phase). After centrifugation at 4°C for 10 min, the cell pellets were frozen immediately in liquid nitrogen. Then, the cells were ground into a powder and dissolved in TRIzol reagent (Invitrogen, Carlsbad, CA). Microarray analysis was performed using Agilent custom 60-mer oligonucleotide microarrays (Shanghai Biochip Co. Ltd., Shanghai, China) as described previously (42, 43). Genes that exhibited greater than 2-fold changes in expression in CAC-smR compared to the expression in the control strain were considered to be differentially expressed.

### Quantitative real-time RT-PCR

For qRT-PCR analyses, RNA was isolated by TRIzol extraction as described previously (42). RNA was reverse transcribed to cDNA using the PrimeScript RT Reagent Kit (TaKaRa, cat. #RR047A). Then, qRT-PCR was carried out in a MyiQ2 two-color real-time PCR detection system (Bio-Rad) with the following conditions: 95°C for 2 min, followed by 40 cycles of 95°C for 15 s, 55°C for 20 s, and 72°C for 20 s. The CAC2679 (42) gene (encoding pullulanase) was used as an internal control. The primers used for qRT-PCR analysis are listed in Table S4.

### Secondary structure analysis and target prediction of sRNA in *C. acetobutylicum*

The secondary structure of sr8384 was generated using PseudoViewer (44). IntaRNA online software (20) was used to screen for the putative sRNA targets across the genome of *C. acetobutylicum* (Target NCBI RefSeq IDs: NC_003030 and NC_001988), in which the 94-nt whole sequence of sr8384 was entered as the query ncRNA.

### In vitro transcription

RNAs were synthesized *in vitro* from PCR-generated DNA fragments using the MEGAscript^TM^ T7 High Yield Transcription Kit (Thermo Fisher Scientific Baltics UAB, Vilnius, Lithuania; AM1334) and then purified using the MEGAclear^TM^ Kit (Thermo Fisher Scientific Baltics UAB, Vilnius, Lithuania; AM1908). The primers used for *in vitro* transcription are listed in Table S4.

### Analysis of RNA-RNA complex formation

RNA-RNA complex formation analysis was performed as previously reported (29). In brief, the 62-nt sr8384 was synthesized and then labeled with Cy5 at the 5’ end. In the RNA hybridization experiment, the Cy5-labeled sr8384 and target RNA were first resolved in TMN buffer (20 mM Tris-acetate (pH 7.5), 2 mM MgCl_2_, 100 mM NaCl) and then incubated at 95°C for 2 min. Next, for proper RNA folding, both the Cy5-labeled sr8384 and target RNA were incubated on ice for 2 min followed by 30 min at 37°C. The Cy5-labeled sr8384 was incubated with various concentrations of target RNA in TMN buffer (containing 0.1 μg/μL tRNA) (Sigma-Aldrich Trading, Shanghai, China; R8508) at 37°C for 15 min. The RNA-RNA complex formation reaction was stopped by adding stop solution (1×TMN, 50% glycerol, 0.5% bromophenol blue and xylene cyanol). Finally, the mixture was loaded onto a 6% native polyacrylamide gel and resolved by electrophoresis (120 V, 4°C, 1 h). The gel was scanned using an FLA-9000 phosphorimager (FujiFilm, Japan) for visualization.

